# Fast and Automated Protein-DNA/RNA Macromolecular Complex Modeling from Cryo-EM Maps

**DOI:** 10.1101/2022.09.29.510189

**Authors:** Andrew Nakamura, Hanze Meng, Minglei Zhao, Fengbin Wang, Jie Hou, Renzhi Cao, Dong Si

## Abstract

Cryo-electron microscopy (cryo-EM) allows a macromolecular structure such as protein-DNA/RNA complexes to be reconstructed in a three-dimensional coulomb potential map. The structural information of these macromolecular complexes forms the foundation for understanding the molecular mechanism including many human diseases. However, the model building of large macromolecular complexes is often difficult and time-consuming. We recently developed DeepTracer-2.0, an artificial intelligence-based pipeline that can build amino acid and nucleic acid backbones from a single cryo-EM map, and even predict the best-fit residues according to the density of side chains. The experiments showed improved accuracy and efficiency when benchmarking the performance on independent experimental maps of protein-DNA/RNA complexes and demonstrated the promising future of macromolecular modeling from cryo-EM maps. Our method and pipeline could benefit researchers worldwide who work in molecular biomedicine and drug discovery, and substantially increase the throughput of the cryo-EM model building. The pipeline has been integrated into the web portal https://deeptracer.uw.edu/.

## Introduction

In 2003, the Worldwide Protein Data Bank^1^ (wwPDB) was formed to ensure Protein Data Bank (PDB) data would be publicly available and archived for researchers to use^2^. In 2012, the resolution revolution in cryo-electron microscopy (cryo-EM) allowed an exponential growth of biological macromolecules structural data, which also extended the protein structure to other types of macromolecules such as RNA/DNA. As of August 24, 2022, there are currently 21,807 published cryo-EM maps. However, only 12,166 structural models are available to these maps^3 4^. Figure 1 shows an overall process of how Cryo-EM data is processed to make a protein-DNA/RNA complex model.

**Fig. 1:**
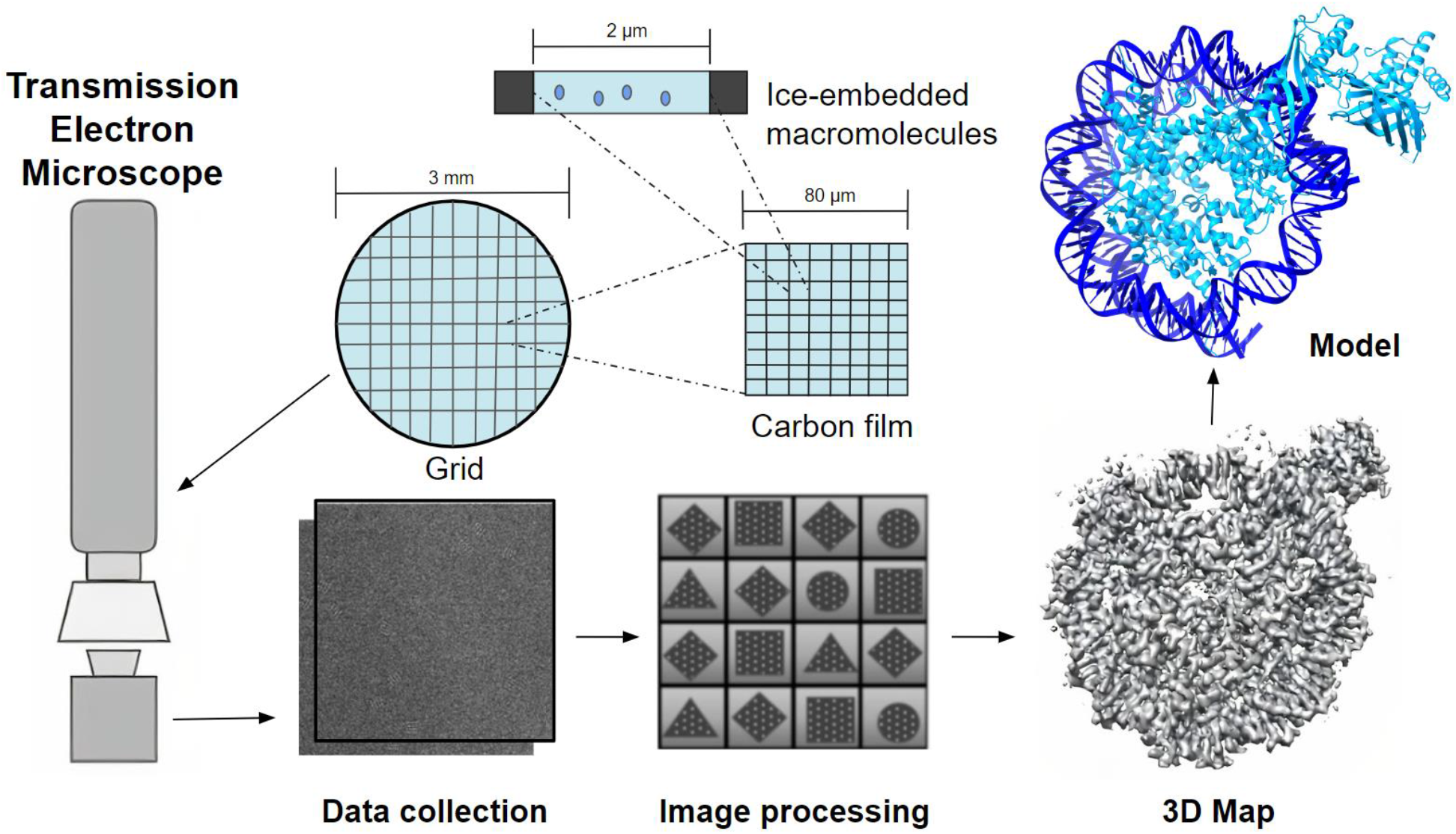
The Overall Cryo-EM Macromolecular Data Processing Pipeline. This process illustrates the use of the microscope to analyze a portion of a biological structure, get the area of interest and transform the image data into a 3D density map. Once complete, the map will contain both the protein-DNA/RNA complex.

DeepTracer is a fully automated deep learning-based method for fast *de novo* multi-chain protein complex structure modeling from cryo-EM maps^5^. When comparing DeepTracer with the state-of-the-art methods of Phenix^4^, Rosetta^6^, and MAINMAST modeling^7^, DeepTracer’s protein carbon alpha (Cα) prediction is more accurate, with higher percent matching averages of Cα at 85-90% and lower root-mean-square deviation (RMSD) values based on the Cα position^5^. However, the previous DeepTracer-1.0 did not account for map regions that involve other macromolecules, such as nucleic acids. This could lead to problems as DNA/RNA, carbohydrates, and fatty acids can potentially be misidentified as amino acids, thereby making an inaccurate prediction with a given cryo-EM map. In this paper, we propose DeepTracer-2.0 to extend the functionality of DeepTracer-1.0 by incorporating the identification of nucleic acids along with amino acids. DeepTracer-2.0 adds segmentation steps for separating cryo-EM maps and a nucleotide U-Net architecture that identifies phosphate and carbon atom positions in the segmented nucleotide cryo-EM map^8^. Combined with preprocessing and postprocessing steps, the DeepTracer-2.0 pipeline achieves fast and accurate macromolecular structure prediction given variable-size cryo-EM inputs. The website, https://deeptracer.uw.edu, allows users to perform automated macromolecular complex modeling using 3D cryo-EM maps and provides users the option to choose between predicting a complex of amino acids and nucleotides, an amino acid only or nucleotide only structure.

### Challenges of Protein-DNA/RNA Macromolecular Modeling

In our early work, we have recognized several key challenges of Protein-DNA/RNA Macromolecular Modeling from cryo-EM maps. This includes separating out the voxels, identifying the critical atoms for each type of macromolecule, and then building the correct chains of macromolecules based on atom positions. It is challenging to distinguish different macromolecules without the accurate separation of the voxels, leading to false positive predictions and the misinterpretation of the voxels as part of the structure.

Amino acids are building blocks that combine to form the protein. Each amino acid has a carbon alpha (Cα) also known as the central carbon, an amino group, and carboxyl group. For each amino acid variation, their R-group defines their characteristic, such as nonpolar, polar acidic, basic, or aromatic. Results are better when the resolution is at 4 angstroms or higher. High resolution has allowed for strategies such as atomic structure modeling, de novo main-chain tracing, structure refinement or a combination of the strategies to identify amino acids^9^. Fasta files are text-based that represent either nucleotide or peptide sequences, in which base pairs or amino acids represent a single-letter code, the description line and sequence are distinguished by a greater-than (“>”) symbol^10^. In fasta files, they are represented as a single letter abbreviation. Although there are 21 different common amino acids in proteins, the previous DeepTracer-1.0 pipeline^5^ focuses on identifying 20 amino acids from the density maps, with selenocystene (SeH), left out of predictions^11^. Each amino acid is attached to another amino acid by a peptide bond, through the carboxyl group and amino group. This resulting chain of amino acids is called a polypeptide chain. Each polypeptide will have a free amino end, the N terminal, as well as a free carboxyl group, the C terminal^12^.

In 2021, about 13% of cryo-EM maps had protein-nucleic acid interactions^9^. Currently, around 3046 released cryo-EM maps with either DNA or RNA included out of 21,806 maps, which brings the map total to 14% of experimental maps^13^. Nucleic acids are macromolecules made up of building blocks called nucleotides. They carry the genetic information of a cell and the instructions for a functioning cell. The two main types of nucleic acids are deoxyribonucleic acid (DNA) and ribonucleic acid (RNA). Each nucleotide consists of three components: a nitrogenous base, a pentose sugar, and a phosphate group. DNA consists of four possible nitrogenous bases, adenine, cytosine, guanine and thymine, while RNA has the same three bases, but has uracil in place of thymine. The nitrogenous base is attached to the 1’ carbon and the phosphate group is attached to the 5’ carbon. Nucleic acids are linear chains of nucleotides, and are held together by phosphodiester linkages between the 3’ carbon nucleotide and 5’ carbon nucleotide phosphate group of another nucleotide. The first nucleotide will have a free phosphate at the end of its 5’ carbon, whereas the last nucleotide will have a free 3’ hydroxyl group at its 3’ carbon^14^. Figure 2 shows the overview of an amino acid and a DNA structure, followed by a closeup of a chain of three amino acids and a chain of three nucleic acids.

**Fig. 2:**
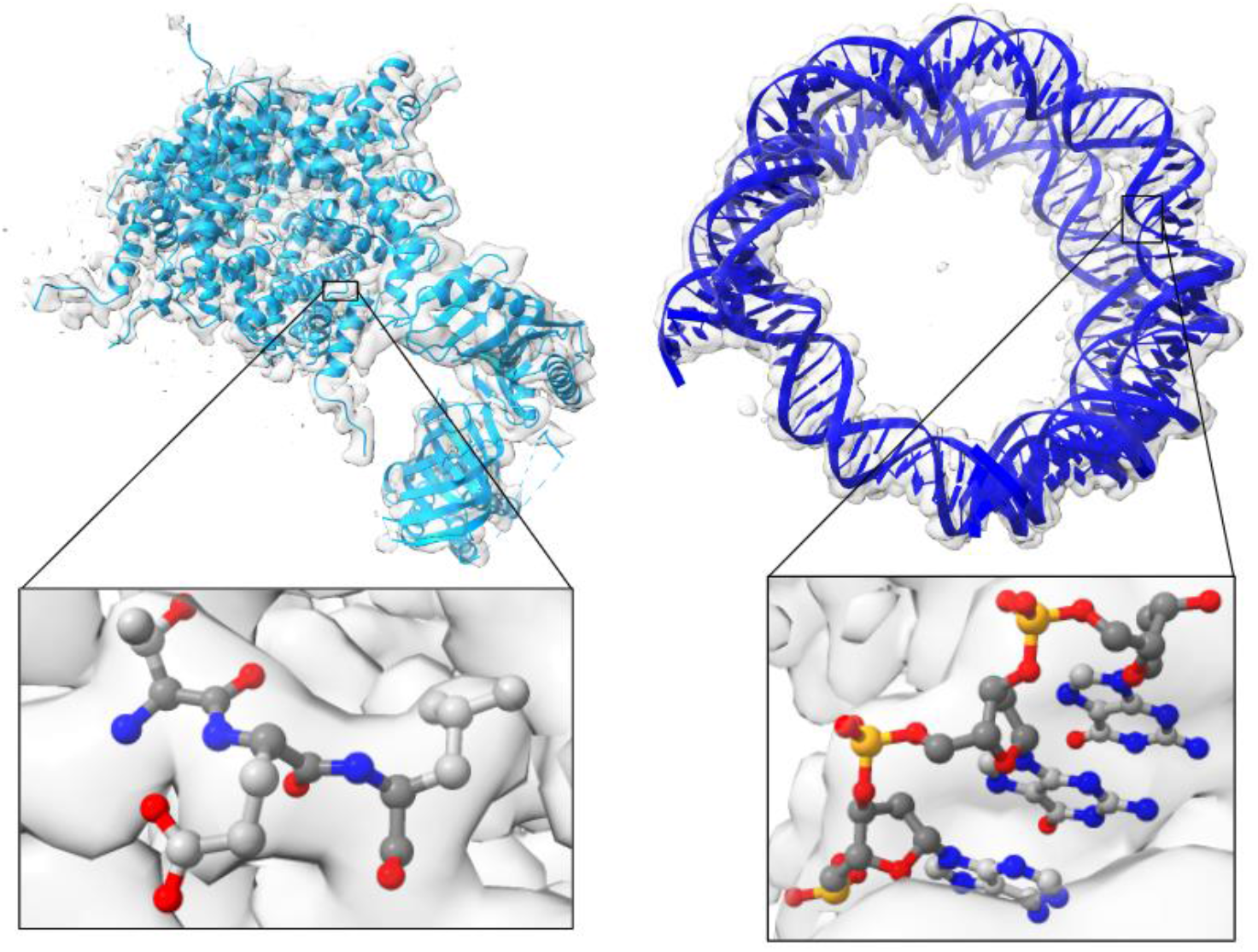
Structural Models of Amino Acids and Nucleotides. The left figure shows a protein structure and a closeup of three amino acids. The right figure shows the nucleic structure of DNA. Blue represents nitrogen atoms, red represents oxygen atoms, orange represents phosphates, light gray shows the side chain carbons, whereas dark gray shows the backbone carbons. Each side chain can have 20 different conformations for amino acids and 5 different kinds of DNA/RNA nucleotides.

The U-Net architecture is a strategy that allows it to work with a moderate number of cryo-EM maps to identify the position of critical atoms for proteins and nucleotides. The U-Net architecture is used in the segmentation steps as well as atom location predictions for amino acids and nucleic acids. The main idea is to supplement a contracting network by successive layers in order to increase the resolution of the output. The high-resolution features from the contracting path are combined with the up-sampled output, and the successive convolutional layer can make a more precise output based on the information. One important modification is upsampling a large number of feature channels. A segmentation map only contains the pixels, for which the full context is available in the input image. Cropping is necessary due to the loss of border pixels in every convolution. It is important to select the input tile size such that all 2×2 max-pooling operations are applied to a layer with an even x- and y-size^15^. The amino acid consists of four U-Nets and the Nucleotide U-Net would consist of two to capture the atom positions.

Once the atoms are identified from their amino acid and nucleotide U-Net architectures, the amino acid inputs and nucleotide inputs proceed to their postprocessing steps. Although the locations of the critical atoms are determined, each macromolecular type requires corresponding steps to connect the backbone atoms correctly. Amino acid and nucleotide chains can have a number of ways to connect each atom to form the backbone, and this makes it impossible to have an exhaustive search of all possible solutions that connect the atoms into chains^5^. Additionally, DeepTracer-2.0 requires a way of respecting the input sequence when building the structural model from cryo-EM maps. Aligning amino acids is challenging due to the fact that some amino acid types have a similar appearance. At lower resolutions, cryo-EM maps lead to the U-Net mismatching the frequency of certain amino acid types^5 16^. Nucleotides have four bases pairs compared to the twenty different amino acid side chain atom types, but still have to correctly distinguish between and pyrimidines as well as distinguish the sugar ribose as DNA or RNA. Therefore, the pipeline adds in nucleotide postprocessing strategies to make the phosphate atoms fit a DNA/RNA structure.

### DeepTracer-2.0 Pipeline

After the nucleotide U-Net was created, all of the U-Net components could be used to separate the macromolecular densities and then predict each macromolecule density as a structure. Our project development uses Python and machine learning algorithms provided by TensorFlow. There are three main processes that allow the prediction of the amino acid and nucleotide macromolecule, seen in Figure 3. The first step is the segmentation to extract the density maps for separate macromolecules. Once separated, the amino acid and nucleotide pipeline work on modeling their structure from their respective density map. When both predictions finish, the protein and DNA/RNA structures can be combined to give a final prediction of the macromolecular model.

**Fig. 3:**
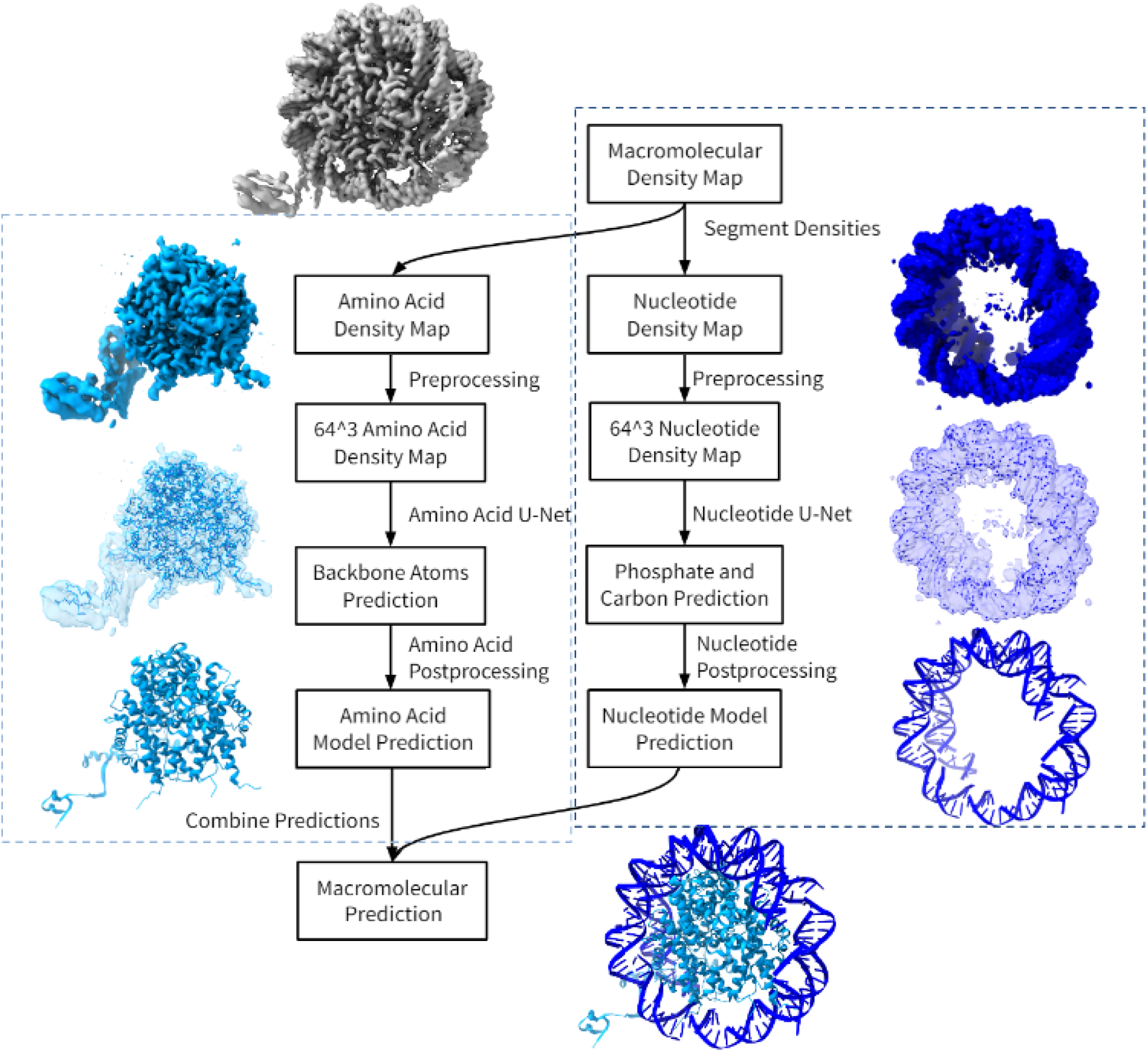
The system design of the Macromolecular Pipeline. The pipeline is organized into four major steps. Segmentation, Preprocessing, their respective machine learning and postprocessing steps. Both amino acids and nucleotides share the same segmentation and preprocessing of their density maps. Once the cryo-EM maps values are normalized and resized to fit a 64^3^ shape, the Amino Acid U-Net determines the positions of the Cα and other backbone atoms, while the Nucleotide U-Net determines the position of phosphate atoms and sugar carbon atoms. Both outputs will be processed through separate postprocessing steps to add on atoms and complete a structure. The models are then combined to generate a complete macromolecular structure. The cyan indicates DeepTracer-1.0’s amino acid pipeline and the dark blue are DeepTracer-2.0’s nucleotide pipeline.

### Convolutional Neural Network and Macromolecular Density Segmentation

Most strategies utilize convolutional neural networks (CNNs) to categorize the voxels^17^. Haruspex utilizes CNN to combine traditional image analysis with machine learning and convolutional filters to obtain high resolution cryo-EM maps that annotate the protein’s secondary structure and DNA/RNA voxel regions. To do this, Haruspex employs a state of the art U-Net architecture to take input that contains the input of 40^3^ voxels segments. The volume is passed through multiple convolutional layers and pooling features which determine the relevant secondary structure elements for proteins or nucleotides. In the second upconvolutional part of the network, activators recover the spatial detail. The output has four channels that annotate the voxel data α-helical, β-strand, nucleotide or unassigned which is used for the segmentation process. Haruspex’s work which reconstructs each cryo-EM density map based on a protein’s secondary structure and DNA/RNA voxel regions^18^. To utilize the output data, the α-helical, β-strand protein and unassigned probabilities are combined to represent the amino acid density of the map. Meanwhile, the nucleotide density probabilities are used to represent the nucleic acid structure. The segmentation network architecture separates density maps that allows each pipeline to model its respective macromolecule in the DeepTracer-2.0 pipeline. Figure 4 shows the result of density segmentation.

**Fig. 4:**
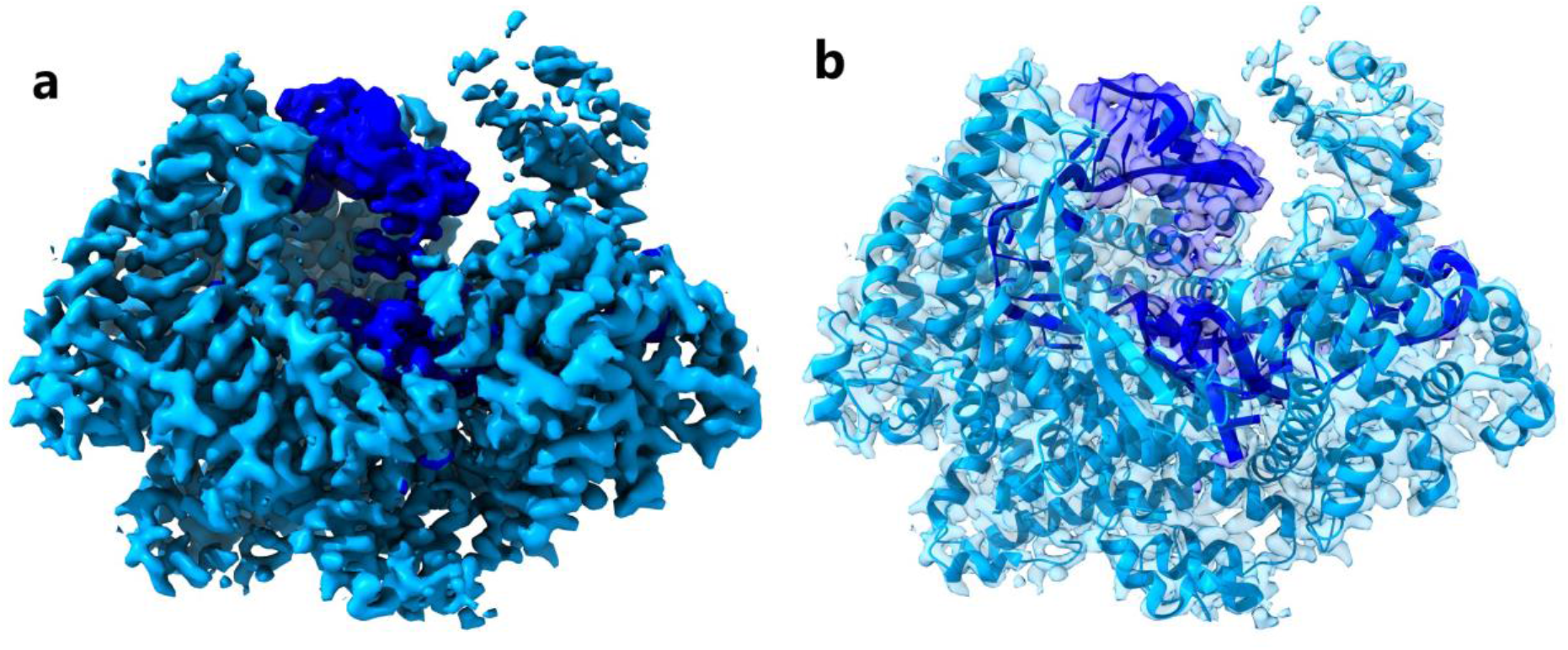
Density Segmentation and Model Structure of EMD-6777. (a) The cyan and dark blue densities are used to solve both the amino acids and nucleic acids structures. On the left is the density map that needs to be separated; the cyan color depicts amino acids where α-helical, β-strand and other data have been combined. Dark blue color depicts RNA density. (b) On the right is the model structure embedded in the density.

### Nucleic Acid Network Architecture

Because DeepTracer focuses on high resolution, only maps with a resolution of 4 Å or better were selected. The initial amino acid network was trained with 1,800 experimental maps and their corresponding deposited model structures^5^. For the nucleotide network training, 293 EMDB/PDB pairs were selected if the cryo-EM map and model represented the same structure and it fit visually well^13^. These maps required at least one protein chain as an α-helix or β-sheet one nucleotide chain that is at least 20 base pairs or larger, and the resolution was 4 Å or better^8^. With these parameters, it narrows down the number of solved structures to a few hundred maps and their respective PDBs.

After separating the densities, each macromolecule would undergo preprocessing to normalize the density value of each voxel from a range of 0 to 1 and resize each map into a 64^3^ voxel size. The preprocessed density map would proceed to the nucleotide U-Net for its atom and backbone prediction. Due to the different molecular structure of nucleotides and amino acids, amino acids use an amino acid U-Net and DNA/RNA use a separate nucleotide U-Net to define the structural aspects of the nucleic acid. Preprocessed cryo-EM maps are fed into the 64^3^ input layer for each U-Net. The atoms for the nucleotide U-Net focus on whether each voxel contains a phosphate atom (P), a carbon one atom (C1’), a carbon four atom (C4’), or no atom. This has a total of four channel outputs. The backbone U-Net determines if each voxel is part of the sugar phosphate backbone, part of the nitrogenous base, or not in either group. This has three different channel outputs. The network architecture for both the atom and backbone U-Nets was heavily focused on determining the structure of the DNA/RNA phosphate backbone. After nucleotide postprocessing is used to refine the phosphate and carbon atom positions, the DeepTracer-2.0 pipeline predicts the nucleotide structure from a cryo-EM map and nucleotide sequence.

### Nucleotide Postprocessing Strategies

Postprocessing steps attempt to reduce the number of phosphates predicted by the U-Net and construct a sugar-phosphate backbone that is consistent with DNA/RNA biological principles. The sugar pucker is predominantly in the C3’ endo (A-DNA or RNA) or a C2’ endo, which corresponds to the DNA’s form (either A or B DNA). Most DNA and RNA conformations fall within the 5.9 Å range as this confirms an A-conformation^19^. However, with larger distances that can appear in B conformations, the postprocessing model allows phosphate atoms that are within 8 Å from each other.

Pseudotorsions were also used to model connecting phosphates to each other. The pseudotorsion simplifies the RNA dihedral angles by utilizing the angles between C1’ atoms and P to distinguish and simplify the construction of the backbone^20^. The presence of the sugar pucker seems to impact the distance between neighboring P atoms.

The Brickworx model then uses the P atoms and Cryo-EM map to finish modeling the nucleotide. Brickworx finds the matching position of double-stranded helical motifs in the cell, and if the structure is RNA, the helical fragments extend to recurrent RNA motifs that can contain single-stranded segments^21^.

### Evaluation Metrics

We examine the results of structure predictions based on their accuracy of amino acid and nucleotide metrics. Testing and training datasets were collected from the publicly available EMDR search tool^13^. DNA samples involving the keywords Repair, Replication, and Splicing, with the filters ‘has DNA’ and ‘< 4Å’. RNA samples had the filters ‘has RNA(no ribosome)’ and ‘< 4Å’ and had no keywords. Ribosomes were avoided for evaluations due to their large size and having nucleotides mixed in with protein density, making ribosomes the hardest samples to predict. From the hundreds of EMDB map entries, the 20 selected cryo-EM maps have a deposited model structure, a fasta sequence, and fall within the resolution of 2 - 4Å with a balance of complexes containing DNA and RNA structures. The metric comparisons are made between Phenix’s pipeline performance and DeepTracer’s pipeline performance. The density map tested has both amino acids and nucleotides, and the nucleotide chains are at least 10 nucleic acids or larger. For map comparisons, no modifications or density map adjustments were made, and were run on the default settings. For Phenix, this was the autosharp and gives the resolution of the density map. Our method compared our metrics with Phenix’s map_to_model in their version 1.19 of the Phenix Suite, using the density maps that are generated from the original density. The metrics comparing the quality of amino acids are RMSD, % matching, % sequence matching and % false positives^22^. The nucleotides metrics are phosphate precision and nucleotide precision. The runtime total was combined for the total length to give an assessment of how long the overall process takes.

For amino acids, accuracy of the atom’s position is measured by the root-mean-square deviation (RMSD) for Cα in amino acid structures^23^. RMSD serves to magnify the significance of errors in the prediction based on the Cα, a lower RMSD value represents a better result. The second is matching percentage; this value represents the proportion of residues from the deposited model that have a matching residue based on the Cα position and is calculated by dividing the total number of residues in the structure. The third metric is a sequence matching percent, which compares if the predicted amino acid has the same type of amino acid. Lastly, the amino acids measure false positives % of predicted residue where no matching residue is found.

Rules for judging nucleotides were taken from Brickworx’s modeling assessment. These general rules were adapted by Gruene and Sheldrick’s *Geometric properties of nucleic acids with potential for autobuilding*^24^. For nucleotides, a P-atom position is considered correct if the distance to the reference structure is within 1.5 Å. The nucleotide position is considered correct if both the P and C1’ atom positions are less than 1.5 and 1.0 Å respectively^21^. Both the phosphate and nucleotides were judged on their precision, which refers to the percentage of predicted structure’s phosphate and nucleotide C1’ atoms positions that are correct.

## Results and Comparisons

The quantitative results for amino acids are displayed in Figure 5, a through d. For the amino acids scatter plots, the average of the map results shows DeepTracer having a lower RMSD, % Matching, % Sequence matching, and % False Positive for the amino acids. Without parameters or manual processing steps, the details on cryo-EM maps are paired with a fasta sequence. In Figure 5, e and f, macromolecules involved in DNA replication had good results. These structures are usually dsDNA on the outside with multiple protein chains coupled on the inside.

**Fig. 5:**
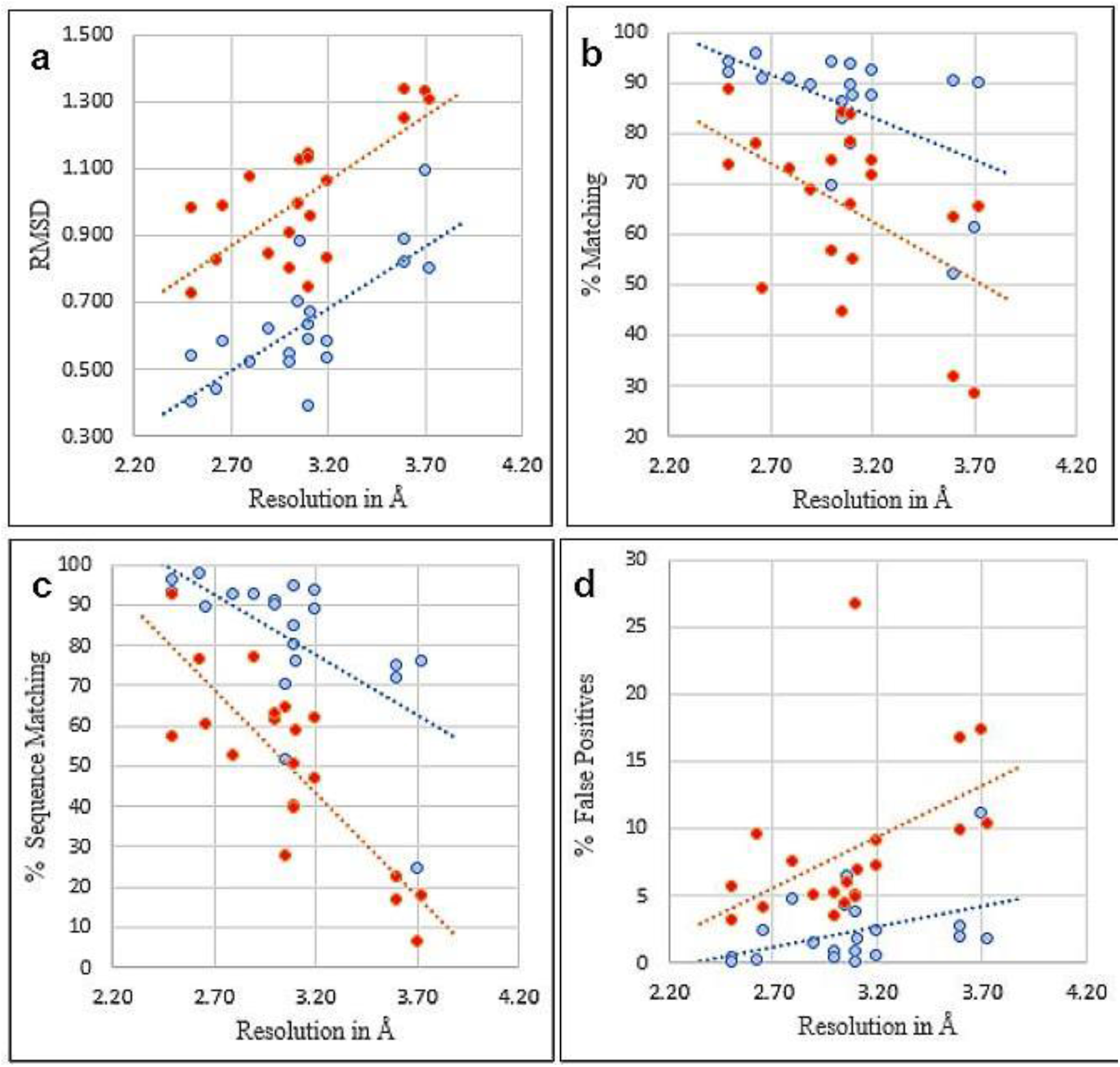
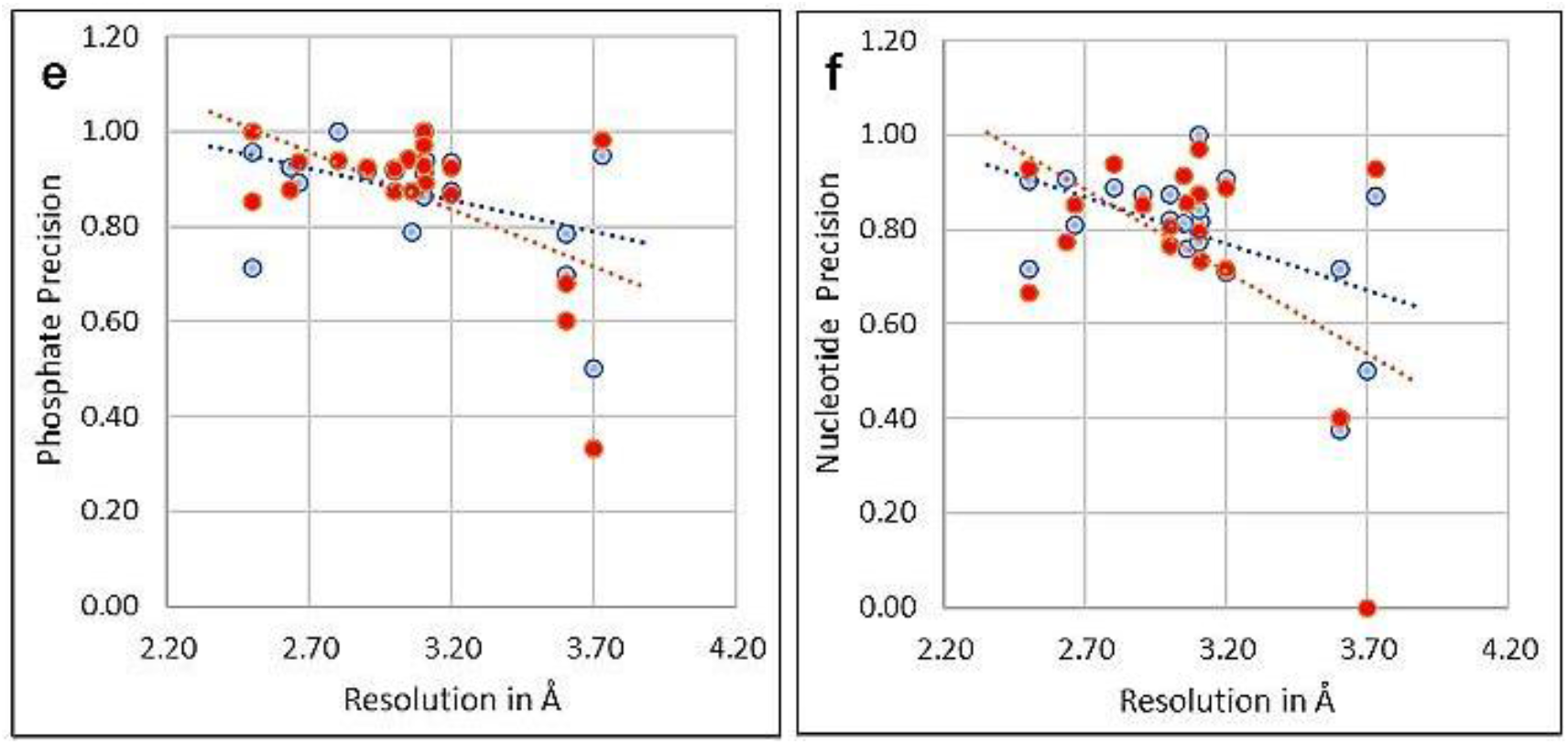
Amino acid and Nucleotide modeling comparison of 20 experimental protein-RNA/DNA cryo-EM maps. DeepTracer models are blue and Phenix models are red. The dotted line represents the trend for each pipeline. (a) RMSD for Cα in amino acid structures. (b) Matching % shows the proportion of residues that have a matching residue, within 3Å. (c) Sequence Matching % shows if the predicted amino acid has the same type of amino acid. (d) False positive % predicted from deposited residues. (e) Nucleotide prediction if the phosphate atom is within 1.5Å. (f) Nucleotide prediction if the phosphate and C1’ atom are within 1.5Å and 1.0Å respectively. For subfigure f, EMD-11550 and EMD-24428 shared the same Nucleotide-CC score (0.4), their phosphate scores were different (0.68 and 0.6).

The amino acid U-Net was capable of performing its prediction as the separated macromolecular densities were tracked well. Notable examples are EMD-6777, EMD-12900 and EMD-31963, which show low RMSD values and great metrics, shown in Figure 6^25 26 27^. These results indicate the capability of DeepTracer’s pipeline to segment the density from the maps and accurately predict the portion of amino acids. Additionally, the nucleotide U-Net was capable of getting a majority of the phosphates required to place the nucleotides in a double-helix structure. EMD-12900 demonstrates a segmented density sample and prediction result with great results. Structures that were nucleosomes, amino acids at the center and nucleotides that surrounded its outside, also have good metrics.

**Fig. 6:**
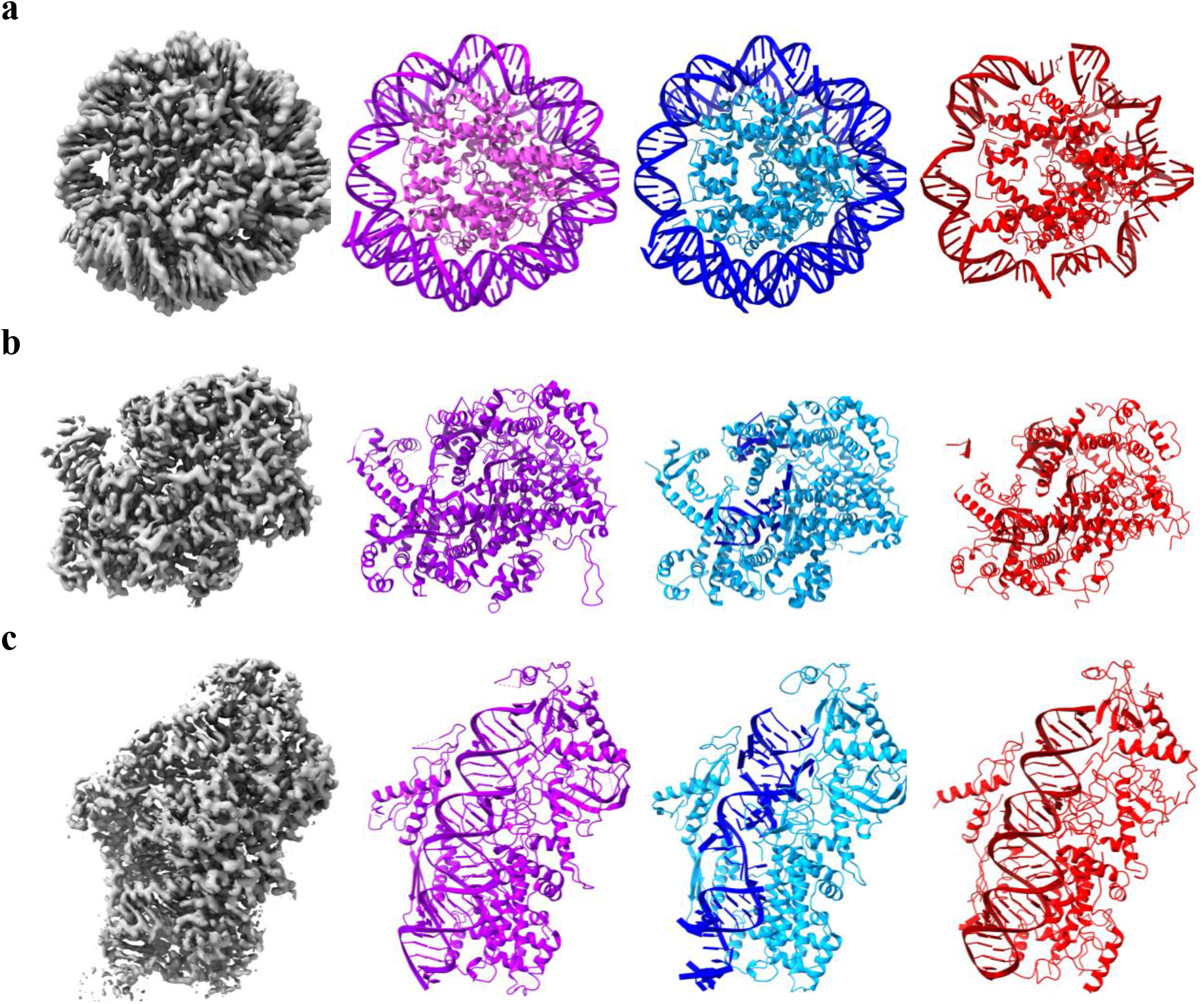
A visual comparison of DeepTracer and Phenix Models from experimental protein-RNA/DNA cryo-EM maps. There are three cryo-EM maps that go through each pipeline in order to predict the structure. Each model built by DeepTracer (blue) and Phenix (red) are compared to each other, while the deposited model (purple) shows how the figure should appear. Darker pigments represent nucleotides whereas lighter pigments represent amino acids. (a) Result of DNA model EMD-12900. Both the DNA and amino acids appear more complete in the DeepTracer pipeline. (b) Result of RNA model EMD-6777. Both maps had issues tracking the single stranded nucleotides. Additionally, Phenix also had a few false positive nucleotides. (c) Result of RNA model EMD-31963. Phenix had a better prediction for the double helix RNA structure, but DeepTracer did better with the amino acid structure.

Figure 7 shows the overall runtime of the pipeline process. DeepTracer’s pipeline was exponentially faster when compared to Phenix’s pipeline in giving a prediction of the complex structure. The smallest structure EMD-25198 took 5 minutes to generate for DeepTracer’s Pipeline, whereas Phenix’s pipeline required 6 hours. For the longer macromolecular complexes, EMD-6941 and EMD-24428, DeepTracer was quickly able to predict both macromolecules taking a bit over 6 minutes while Phenix required over a day to predict the overall structure.

**Fig. 7:**
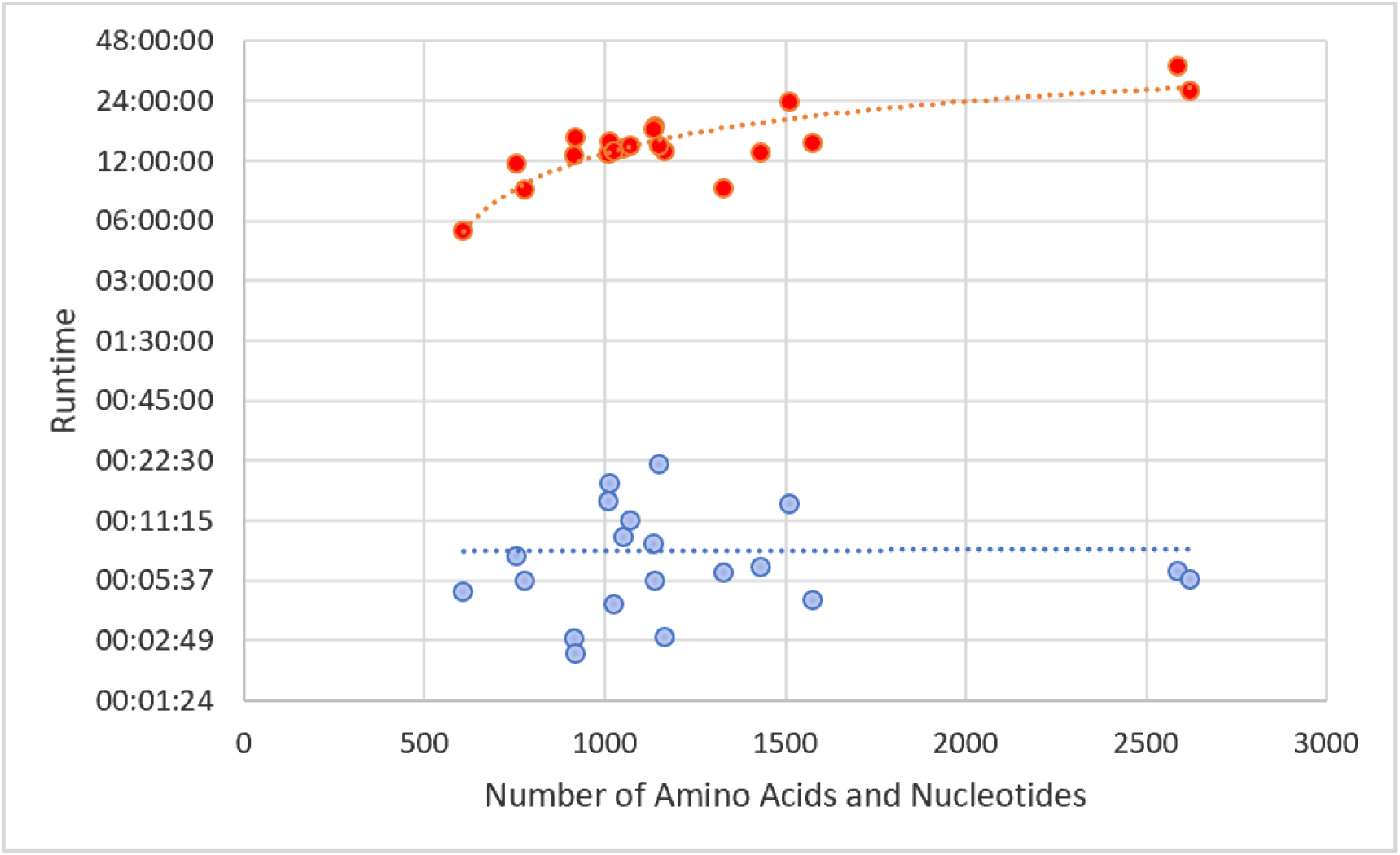
Computational runtime comparison of 20 experimental protein-RNA/DNA maps. Dotted lines represent the trend for each method. The times are shown on a logarithmic scale and combine the runtime totals of amino acid and nucleotide methods. DeepTracer models are blue and Phenix models are red.

The summary of comparison between DeepTracer and Phenix is provided in Table 1 and details of the comparison can be found in the Supplementary Tables. The amino acid and nucleotide density that is used for each method comes from the original density map, with each map resolution in the range of 2.0 - 4.0Å. Each program uses its methods of using the density to perform its evaluation of the macromolecule. Each pipeline lists the average values for the total 20 maps tested. The 2nd to 6th columns of the table show amino acids metrics and the last two columns show nucleotide metrics. The results for each individual density map can be found on the supplementary page.

**Table 1:** Summary of comparison between DeepTracer and Phenix.

| Pipeline | RMSD | % Matching | % Sequence Matching | % False Positives | % False Connections | Phosphate Precision | Nucleotide Precision |
| --- | --- | --- | --- | --- | --- | --- | --- |
| DeepTracer | 0.638 | 85.32 | 81.49 | 2.467 | 1.848 | 0.872 | 0.793 |
| Phenix | 1.017 | 65.44 | 49.71 | 8.416 | 4.102 | 0.866 | 0.752 |

## Discussion

With the new implementation of the macromolecular density segmentation and nucleic acid modeling, DeepTracer-2.0 pipeline is capable of predicting protein-DNA/RNA macromolecular complexes from the cryo-EM maps. The concept of accurate density segmentation allows for researchers to submit cryo-EM maps that can contain different macromolecules in order to identify the types of macromolecules involved in each voxel. As shown by previous results, the amino acids were able to have low RMSD and DNA/RNA contained a high phosphate precision. It is also observed that low map quality could lead to suboptimal segmentation and modeling results. Also, some large maps could increase the computational time. For future work, other deep learning and artificial intelligence methods could be explored to train on the accumulated amino acid density map and nucleic acid density map data and target on single strand RNA as well as larger complexes. In addition, refining the sugar phosphate backbone to model the secondary structure of DNA/RNA could lead to more accurate models.

