## Supplementary Materials for "Fast and Automated Protein-DNA/RNA Macromolecular Complex Modeling from Cryo-EM Maps"

**Supplementary Methods**

The EMDataResource has both density maps and deposited model structures that serve as the ground truth when training the U-Net Networks. In order to design the nucleotide U-Net, we avoided simulated data and used maps that contain both macromolecules. Maps having a resolution of 4 angstroms (Å) or better were prioritized. Initially, a total of 900 experimental maps containing both amino acids and nucleotides (DNA/RNA) had been downloaded, which contained their density map and solved structures. After visually examining the maps through Chimera, it was evident that despite being listed as 4 Å, the overall map quality was low and other maps containing a minimal number of nucleotides or unusual structures were not suitable for training data.

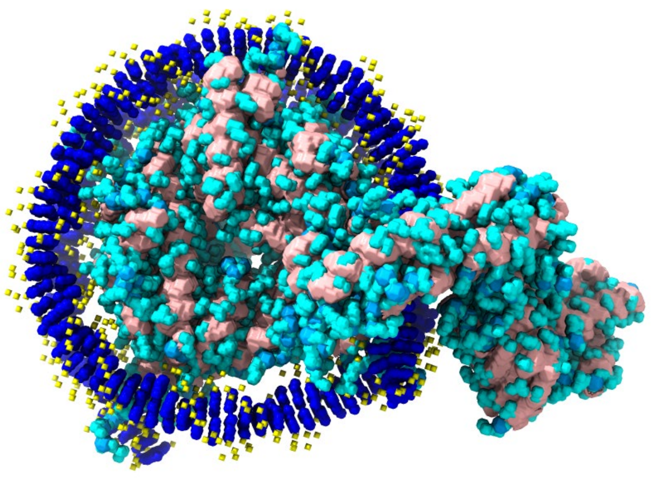

**SI Fig. 1: EMD-12897 Map of Amino Acid and Nucleotides**. The blue and yellow densities represent the atoms mask and dark blue are the backbone mask for nucleotides. Pink densities represent the secondary structure of the amino acid masks, followed by a cyan for the backbone mask and blue for the amino acid mask.

To balance out the data, another search was redefined in biological functions. This included maps from DNA Repair, DNA replication, spliceosomes and RNA samples with no ribosomes. These datasets selected would also have high resolution and contain both protein and

DNA/RNA. Additionally, to avoid redundant density data maps, only one density map from a group of similar maps was chosen if conformation was extremely similar and only varied by a slight difference in its protein content, nucleic acid content or resolution. After screening from these datasets, a total of 163 density maps with resolutions from 2.0 – 3.9Å were selected to make up the DNA/RNA U-Net. In order to use the nucleotide density, the 163 density maps were segmented into DNA/RNA densities. This step includes segmenting the solved PDB models for each density map. Preprocessing of nucleotide density maps normalizes the density values between 0 and 1. Two nucleotide masks, an atom and a backbone mask are generated from each nucleotide density map. In training the maps, the set was randomly split into training and validation sets with an 80:20 ratio.

For the amino acid U-Net, we utilized the same training set used for Deep-Tracer-1.0. The two figures represent how the U-Nets are organized.

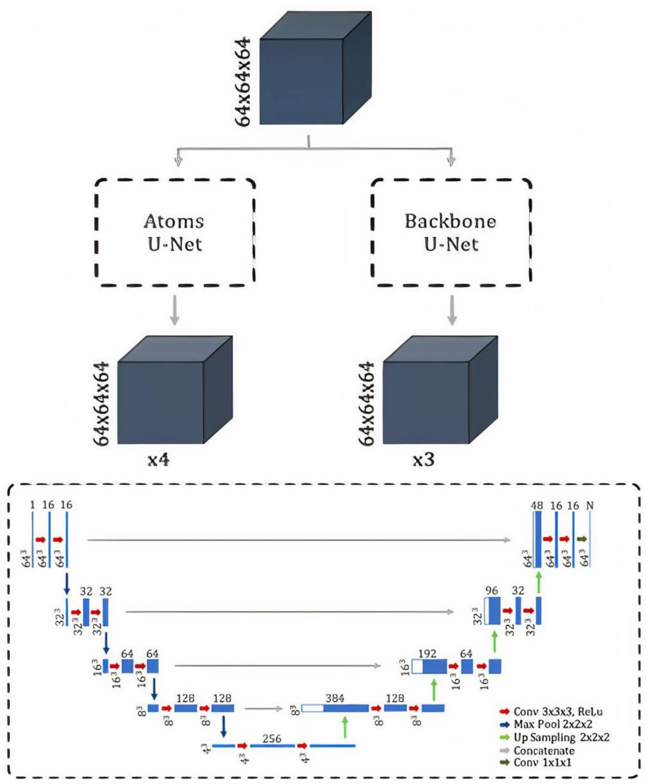


**SI Fig. 2: DeepTracer's overview of the nucleotide neural network**. The top shows two parallel U-Nets, with gray boxes that show the input and output of their maps. This is similar to the amino acid neural network, except the atoms that are mapped are the P, C1' and C4' when compared to Cα, N and C atoms. The bottom dashed box details the architecture of each parallel U-Net.

**Specifications**

For DeepTracer training, pipeline test, and Phenix pipeline testing, the tests were performed on the DAIS2 machine. For specifications, DAIS2 has CPU uses an Intel Xeon CPU E5-2623 v2 @ 2.6 Ghz, 1 socket, core per socket: 4, thread per core: 2, a total of 8 threads. When the GPU was applicable, the graphics card was Nvidia-SMI 418.87.00, the CUDA version 10.1 two GeForce RTX 1080 ti and one Titan RTX. There was a total 128 GB of ram available for both pipelines. For the DeepTracer pipeline and the phenix pipeline, the python version is 3.8.13 and tensorflow was 2.3.2, using a 10.1 cuda toolkit.

### Supplementary Tables

**SI Table 1: Amino Acid Comparison with DeepTracer's Pipeline and Phenix's Map_to_Model Output.** The amino acid density that is used for each method comes from the original density map. Each program uses its methods to separate the density and perform its evaluation of the macromolecule. At the end of the table, an average is listed for each pipeline.

| DeepTracer | Resolution (Å) | Predicted Residues | Model Residues | RMSD | % Matching | % Sequence Matching | % False Positives | % False Connections |
|---|---|---|---|---|---|---|---|---|
| 6777 | 3.2 | 1039 | 1117 | 0.579 | 92.48 | 93.51 | 0.577 | 0.199 |
| 6941 | 3.0 | 2375 | 2509 | 0.548 | 93.90 | 91.17 | 0.800 | 0.825 |
| 8945 | 2.63 | 1171 | 1220 | 0.438 | 95.82 | 97.52 | 0.171 | 0.174 |
| 10069 | 3.2 | 763 | 852 | 0.530 | 87.44 | 88.99 | 2.359 | 0.972 |
| 11550 | 3.6 | 1396 | 1520 | 0.888 | 90.13 | 74.96 | 1.862 | 2.982 |
| 11601 | 3.11 | 951 | 1070 | 0.672 | 87.38 | 75.72 | 1.682 | 2.275 |
| 11692 | 2.5 | 1031 | 1120 | 0.537 | 91.79 | 93.19 | 0.291 | 0.801 |
| 11993 | 3.1 | 803 | 991 | 0.588 | 78.00 | 80.08 | 3.736 | 0.963 |
| 11995 | 2.8 | 945 | 991 | 0.522 | 90.92 | 92.79 | 4.656 | 0.229 |
| 12827 | 2.66 | 653 | 703 | 0.585 | 90.61 | 89.48 | 2.450 | 0.968 |
| 12897 | 3 | 798 | 1144 | 0.520 | 69.49 | 89.81 | 0.376 | 1.057 |
| 12900 | 2.5 | 717 | 764 | 0.403 | 93.85 | 96.09 | 0.000 | 1.563 |
| 23600 | 3.05 | 602 | 669 | 0.702 | 86.10 | 70.14 | 4.319 | 3.279 |
| 24428 | 3.6 | 1364 | 2551 | 0.822 | 52.06 | 71.76 | 2.639 | 3.073 |
| 25198 | 3.7 | 385 | 558 | 1.094 | 61.29 | 24.56 | 11.169 | 8.156 |
| 30767 | 3.06 | 829 | 937 | 0.880 | 82.82 | 51.55 | 6.393 | 5.035 |
| 31106 | 3.1 | 698 | 779 | 0.392 | 89.60 | 94.84 | 1.828 | 0.000 |
| 31963 | 3.1 | 797 | 845 | 0.630 | 93.49 | 84.94 | 0.878 | 0.925 |
| 31964 | 3.73 | 786 | 860 | 0.801 | 89.77 | 76.17 | 1.781 | 2.877 |
| 32389 | 2.90 | 871 | 960 | 0.619 | 89.48 | 92.43 | 1.378 | 0.601 |
| **Averages** | | | | **0.638** | **85.32** | **81.49** | **2.467** | **1.848** |
| Phenix | Resolution (Å) | Predicted Residues | Model Residues | RMSD | % Matching | % Sequence Matching | % False Positives | % False Connections |
| 6777 | 3.2 | 898 | 1117 | 1.064 | 74.57 | 46.94 | 7.24 | 3.49 |
| 6941 | 3.0 | 1935 | 2509 | 0.906 | 74.45 | 61.30 | 3.46 | 2.46 |

| | | | | | | | | |
|---|---|---|---|---|---|---|---|---|
| 8945 | 2.63 | 1053 | 1220 | 0.825 | 78.03 | 76.58 | 9.59 | 1.89 |
| 10069 | 3.2 | 672 | 852 | 0.829 | 71.71 | 61.70 | 9.08 | 1.90 |
| 11550 | 3.6 | 1158 | 1520 | 1.251 | 63.42 | 22.61 | 16.75 | 9.96 |
| 11601 | 3.11 | 634 | 1070 | 0.953 | 55.14 | 58.98 | 6.94 | 2.72 |
| 11692 | 2.5 | 875 | 1120 | 0.979 | 73.66 | 57.21 | 5.71 | 4.06 |
| 11993 | 3.1 | 686 | 991 | 1.143 | 65.79 | 40.03 | 4.96 | 4.13 |
| 11995 | 2.8 | 779 | 991 | 1.072 | 72.75 | 52.43 | 7.45 | 5.12 |
| 12827 | 2.66 | 361 | 703 | 0.987 | 49.22 | 60.12 | 4.16 | 3.80 |
| 12897 | 3 | 685 | 1144 | 0.801 | 56.73 | 62.87 | 5.26 | 2.40 |
| 12900 | 2.5 | 699 | 764 | 0.726 | 88.61 | 92.76 | 3.15 | 2.45 |
| 23600 | 3.05 | 589 | 669 | 0.993 | 84.16 | 64.30 | 4.41 | 2.46 |
| 24428 | 3.6 | 893 | 2551 | 1.338 | 31.56 | 17.02 | 9.85 | 4.70 |
| 25198 | 3.7 | 191 | 558 | 1.332 | 28.32 | 6.33 | 17.28 | 10.08 |
| 30767 | 3.06 | 447 | 937 | 1.123 | 44.82 | 27.86 | 6.04 | 4.22 |
| 31106 | 3.1 | 888 | 779 | 0.746 | 83.57 | 50.38 | 26.69 | 1.43 |
| 31963 | 3.1 | 694 | 845 | 1.129 | 78.11 | 39.55 | 4.90 | 5.08 |
| 31964 | 3.73 | 626 | 860 | 1.306 | 65.23 | 18.00 | 10.38 | 6.67 |
| 32389 | 2.90 | 696 | 960 | 0.845 | 68.85 | 77.16 | 5.03 | 3.02 |
| **Averages** | | | | **1.017** | **65.44** | **49.71** | **8.416** | **4.102** |

**SI Table 2: Nucleotide Comparisons with *DeepTracer*'s Pipeline and Phenix's Map to Model Outputs**. The nucleotide density that is used for obtaining these results comes from the original density map.

| DeepTracer | Resolution (Å) | Predicted Nucleotides | Model Nucleotides | Phosphate Precision | Nucleotide Precision | Nucleotide Type |
|---|---|---|---|---|---|---|
| 6777 | 3.2 | 27 | 51 | 0.875 | 0.708 | RNA |
| 6941 | 3.0 | 72 | 81 | 0.922 | 0.875 | DNA |
| 8945 | 2.63 | 234 | 292 | 0.924 | 0.906 | DNA |
| 10069 | 3.2 | 255 | 300 | 0.934 | 0.905 | DNA |
| 11550 | 3.6 | 46 | 56 | 0.700 | 0.375 | DNA |
| 11601 | 3.11 | 225 | 262 | 0.939 | 0.818 | DNA |
| 11692 | 2.5 | 17 | 21 | 0.714 | 0.714 | DNA |
| 11993 | 3.1 | 43 | 21 | 1.000 | 1.000 | RNA |
| 11995 | 2.8 | 37 | 22 | 1.000 | 0.889 | RNA |
| 12827 | 2.66 | 46 | 78 | 0.892 | 0.811 | DNA/RNA |
| 12897 | 3 | 229 | 290 | 0.916 | 0.819 | DNA |
| 12900 | 2.5 | 270 | 289 | 0.955 | 0.902 | DNA |
| 23600 | 3.05 | 64 | 88 | 0.943 | 0.811 | DNA/RNA |

| | | | | | | |
|---|---|---|---|---|---|---|
| 24428 | 3.6 | 14 | 71 | 0.786 | 0.714 | RNA |
| 25198 | 3.7 | 7 | 50 | 0.500 | 0.500 | RNA |
| 30767 | 3.06 | 44 | 88 | 0.788 | 0.758 | RNA |
| 31106 | 3.1 | 236 | 292 | 0.914 | 0.842 | DNA |
| 31963 | 3.1 | 70 | 71 | 0.864 | 0.773 | RNA |
| 31964 | 3.73 | 45 | 61 | 0.949 | 0.872 | RNA |
| 32389 | 2.90 | 96 | 175 | 0.919 | 0.872 | DNA/RNA |
| **Averages** | | | | **0.872** | **0.793** | |

| Phenix | Resolution (Å) | Predicted Nucleotides | Model Nucleotides | Phosphate Precision | Nucleotide Precision | Nucleotide Type |
|---|---|---|---|---|---|---|
| 6777 | 3.2 | 37 | 51 | 0.926 | 0.889 | RNA |
| 6941 | 3.0 | 65 | 81 | 0.919 | 0.806 | DNA |
| 8945 | 2.63 | 184 | 292 | 0.877 | 0.772 | DNA |
| 10069 | 3.2 | 171 | 300 | 0.868 | 0.717 | DNA |
| 11550 | 3.6 | 12 | 56 | 0.600 | 0.400 | DNA |
| 11601 | 3.11 | 105 | 262 | 0.892 | 0.735 | DNA |
| 11692 | 2.5 | 19 | 21 | 1.000 | 0.929 | DNA |
| 11993 | 3.1 | 25 | 21 | 1.000 | 0.875 | RNA |
| 11995 | 2.8 | 30 | 22 | 0.938 | 0.938 | RNA |
| 12827 | 2.66 | 48 | 78 | 0.936 | 0.851 | DNA/RNA |
| 12897 | 3 | 153 | 290 | 0.872 | 0.765 | DNA |
| 12900 | 2.5 | 136 | 289 | 0.851 | 0.664 | DNA |
| 23600 | 3.05 | 76 | 88 | 0.941 | 0.912 | DNA/RNA |
| 24428 | 3.6 | 146 | 71 | 0.680 | 0.400 | RNA |
| 25198 | 3.7 | 65 | 50 | 0.333 | 0.000 | RNA |
| 30767 | 3.06 | 100 | 88 | 0.873 | 0.855 | RNA |
| 31106 | 3.1 | 156 | 292 | 0.929 | 0.795 | DNA |
| 31963 | 3.1 | 71 | 71 | 0.971 | 0.971 | RNA |
| 31964 | 3.73 | 72 | 61 | 0.981 | 0.926 | RNA |
| 32389 | 2.90 | 125 | 175 | 0.926 | 0.851 | DNA/RNA |
| **Averages** | | | | **0.866** | **0.752** | |
